# Reviving the Desired Gains Index: An optimal solution for parent selection in public plant breeding programs

**DOI:** 10.1101/2024.07.21.603926

**Authors:** C.R. Werner, K.A. Gardner, D.J. Tolhurst

## Abstract

The Desired Gains Index is an optimal solution for parent selection in public plant breeding programs. It enables breeders to quantify their breeding objectives in terms of desired genetic gains and facilitates the efficient simultaneous improvement of multiple quantitative traits in a breeding population without the need for economic weights. We deliberately chose the term “optimal” here, which is typically associated with the profit-oriented selection indices commonly used in animal breeding, such as the Smith-Hazel Index. Our intention is to refute the perception that the Desired Gains Index is less efficient than the Smith-Hazel Index since both approaches maximise expected genetic gains in proportion to the breeding objective. To achieve this, we first review the relationship between the Desired Gains Index and the Smith-Hazel Index to show that desired gains are actually a form of economic weighting expressed as an improvement ratio for the traits under selection. We then present a general form of the Desired Gains Index to leverage best linear unbiased prediction (BLUP), which enables seamless integration of pedigree or genomic relationship information (e.g., genomic estimated breeding values) between the selection candidates and other related individuals. Finally, using stochastic simulation, we compare the performance of different parent selection strategies, including index selection, independent culling, and selection of extreme genotypes. The objective of these demonstrations is to convey the potential impact and benefits of the Desired Gains Index to a broader audience without the need for a deep understanding of selection theory and the equations presented here.

## 1 Introduction

We advocate for the Desired Gains Index as an optimal solution for parent selection in public plant breeding programs. The Desired Gains Index enables breeders to quantify their breeding objectives in terms of desired genetic gains (i.e., desired response to selection) and facilitates the efficient simultaneous improvement of multiple quantitative traits in a breeding population without the need for economic weights.

Plant breeding programs aim to simultaneously improve multiple traits of commercial importance to release varieties that meet the complex and dynamic requirements of growers, processors, and end-users (Covarrubia Pazaran et al., 2022). In addition to superior yield under the impact of various biotic and abiotic stresses, novel varieties must also meet minimum standards for several quality and agronomic traits, such as protein and oil content, plant height, and flowering time (Bernardo, 2010). However, simultaneous improvement of multiple traits is challenging. Traits are often unfavourably correlated (e.g., Erskine et al., 1985; Kwon and Torrie, 1964; Meredith and Bridge, 1971; Triboi et al., 2006), and selection on one trait affects the response to selection that can be realised in other traits (Falconer and Mackay, 1995).

Index selection is generally the most efficient method to improve a population for multiple quantitative traits simultaneously (Young, 1961; Finney, 1962; Henderson, 1963; Pešek and Baker, 1969a; Batista et al., 2021). The traits of interest are first weighted by their importance in terms of the breeding objective and then combined into a single value, representing an individual’s merit. The breeding objective can be defined, for example, in terms of total economic profit, gain per trait, or some combination of both (Brascamp, 1984). The individuals with the highest selection index are the best parents to achieve the breeding objective in the next generation.

The use of index selection for simultaneous improvement of multiple traits in plant breeding was first proposed by H. Fairfield Smith in 1936. Smith introduced the concept that the “genotypic worth of a plant” as a parent could be expressed as a linear function of the genotypic values of all traits of interest (Baker, 1986). A few years later, Hazel and Lush (1942) and Hazel (1943) proposed the idea of a selection index in animal breeding (also see Hazel et al., 1994).

Smith, Hazel, and Lush formulated their selection index theory exclusively from an economic perspective. What Smith described as the “genotypic worth of a plant” is now commonly referred to as its aggregate genotype (or aggregate genetic value/merit), a function of the breeding values for the traits of interest, weighted by their economic value. The aggregate genotype of an individual can be written as:

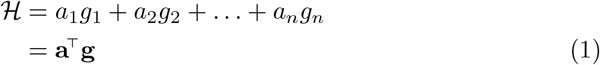

where **a** is the vector of economic weights for the *n* traits and **g** is the vector of multivariate breeding values.

Hazel (1943) defined the economic weight for a trait as “the expected independent monetary return from a unit of change”, that is, the economic value of improving the trait when all other traits remain unchanged. Assuming the economic weights of each trait are known without error, the individuals with the highest aggregate genotype are the parents that will maximise profit in the next generation.

In practice, however, the true breeding values in Eq. 1 are unknown, and the aggregate genotype must be estimated. Smith (1936) stated that this is best achieved using a linear function of the phenotypes, which are combined into a selection index. The approach is commonly known as the “Smith-Hazel Index”. The Smith-Hazel Index for an individual is calculated as:

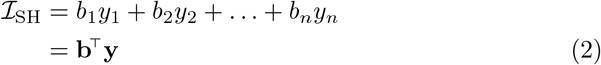

where **b** is the vector of selection index coefficients for the *n* traits which are a function of the economic weights in **a** and **y** is the vector of phenotypes. The Smith-Hazel Index, ℐ_SH_, is a correlated trait with the aggregate genotype, ℋ. Smith (1936) and Hazel (1943) showed that selection on ℐ_SH_ maximises the correlation between ℐ_SH_ and ℋ, and, therefore, maximises the response to selection in ℋ when the vector of index coefficients in Eq. 2 is estimated as:

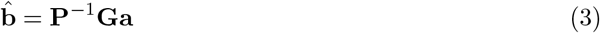

where **P** is the phenotypic variance-covariance matrix for the *n* traits, **G** is the additive genetic variance-covariance matrix, and **a** is the vector of economic weights.

The vector of index coefficients in Eq. 3 gives the Smith-Hazel Index for an individual:

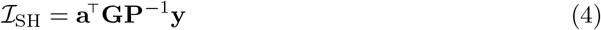

which is a linear function of 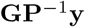. The term **GP**^−1^ can be viewed as multivariate narrow-sense heritability, based on the population parameters **G** and **P**. It facilitates the exchange of information between correlated traits and, when multiplied with **y**, shrinks the phenotypes according to their heritability. That is, **GP**^−1^ converts the phenotypes in **y** into multivariate best linear unbiased predictors (BLUPs) of the breeding values, given by 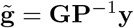 and often referred to as estimated breeding values (EBVs). Assuming that **G** and **P** are known without error, the vector of index coefficients yields what is generally considered the optimal index ℐ_SH_ (Baker, 1986).

Henderson (1951, as cited in Hazel et al., 1994) later showed that the Smith-Hazel Index could be calculated using EBVs directly as input values instead of phenotypes. In this case, the EBVs are obtained from fitting a linear mixed model to **y** and solving Henderson’s mixed model equations for 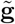 (Henderson, 1963). The Smith-Hazel Index in Eq. 4 can, therefore, be rewritten as:

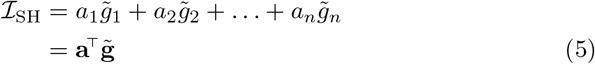

where **a** is the vector of economic weights for the *n* traits and 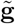 is the vector of multivariate EBVs. This approach has the advantage that the EBVs can be adjusted for varying amounts of information for each individual (Henderson, 1963, 1973). Furthermore, it enables seamless integration with genomic selection, allowing the utilisation of genomic estimated breeding values (GEBVs).

Despite the routine adoption of the Smith-Hazel Index and its various modifications to maximise economic profit in animal breeding programs (Brascamp, 1984), index selection remains a widely underutilised tool in public plant breeding programs. The derivation of economic weights is a complex, laborious, and costly process, and public plant breeding programs often lack the resources to accurately quantify traits in terms of profitability.

Pešek and Baker (1969b) recognised that “few breeders are prepared to assign relative economic weights to traits but most would be willing to specify the amount of gain they would like to see in each trait in a given selection program”. Half a century later, their statement is still remarkably relevant. To overcome the need for economic weights, Pešek and Baker proposed a modification of the Smith-Hazel Index in which the expected response to index selection was replaced by a vector of desired responses (i.e., desired genetic gains). Solving for the vector of index coefficients, they obtained:

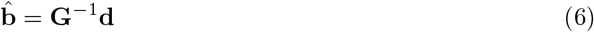

where **G** is the additive genetic variance-covariance matrix for the *n* traits and **d** is the vector of desired genetic gains. Redefining the index coefficients in this manner enables breeders to express their breeding objective in terms of desired genetic gains with reference to the current population mean rather than profit. The vector of index coefficients in Eq. 6 gives the Desired Gains Index for an individual:

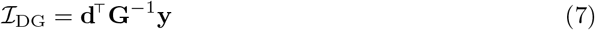

which maximises the expected response to index selection, i.e., realised genetic gains, in proportion to the desired gains in **d** (Pešek and Baker, 1969b). Furthermore, this approach simplifies stabilising key agronomic traits, such as plant height or flowering time, by simply setting their desired gains to zero. In fact, the desired gains represent a form of economic weighting expressed as an improvement ratio for the traits in **y**.

We advocate for the Desired Gains Index as an optimal solution for parent selection in public plant breeding programs. The Desired Gains Index facilitates the efficient simultaneous improvement of multiple quantitative traits in a breeding population without the need for economic weights, which are often unknown and difficult to obtain. Breeders generally find the concept of desired gains highly intuitive because it can be directly aligned with their objective of developing improved crop varieties.

We deliberately chose the term “optimal” here, which is typically only associated with the profit-oriented selection indices used in animal breeding. Our intention is to refute the widespread perception that the Desired Gains Index is less efficient than the Smith-Hazel Index, which is only strictly true when the breeding objective is profit maximisation rather than maximising genetic gain and the economic weights are known without error.

The paper consists of three parts. In the first part, we review the relationship between the Desired Gains Index and the Smith-Hazel Index. Based on Brascamp (1984), we show that the desired gains in **d** can be expressed in terms of the economic weights in **a**, thereby providing evidence that the Desired Gains Index is not per se less efficient than the Smith-Hazel Index. In the second part, we derive an equation for the Desired Gains Index to leverage Best Linear Unbiased Prediction (BLUP), similar to the approach introduced by Henderson (1951, as cited in Hazel et al., 1994) for the Smith-Hazel Index. In the third part, using stochastic simulation, we compare the performance of different parent selection strategies, including index selection for population improvement, independent culling, and selection of extreme genotypes. While the insights from these simulations are in principle not new, we believe that our demonstrations will help to convey the potential impact and benefits of the Desired Gains Index to a broader audience without the need for a deep understanding of selection theory and the equations presented in this paper.

## 2 Materials and methods

The Methods section consists of two parts. In the first part, we propose a modification of the Desired Gains Index that incorporates BLUPs instead of phenotypes or adjusted phenotypes. Our derivation utilises the relationship between the Smith-Hazel Index and the Desired Gains Index as introduced by Pešek and Baker (1969b). In the second part, we describe the stochastic simulations used to demonstrate the advantages of using a Desired Gains Index for parental selection compared to other selection strategies. Specifically, we compare its efficiency to that of the Smith-Hazel Index, independent culling (also called truncation selection), and selection of extreme genotypes, for equally improving two quantitative traits within a breeding population while stabilising a third quantitative trait.

### 2.1 Reviving the Desired Gains Index

#### The relationship between the Desired Gains Index and the Smith-Hazel Index

When Pešek and Baker (1969b) introduced the concept of a Desired Gains Index, they highlighted that desired gains are essentially a form of economic weights. The direct relationship between the desired gains in **d** and the economic weights in **a**, however, remained implicit until Brascamp (1984) showed how to obtain the economic weights that can achieve genetic gains equal to those targeted by the desired gains. A similar approach is adopted in the following to demonstrate how to identify the form of **d** and **a** that generate identical index coefficients for the Desired Gains Index in Eq. 7 and the Smith-Hazel Index in Eq. 4, denoted by 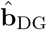 and 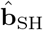, respectively. The vectors of index coefficients are given by:

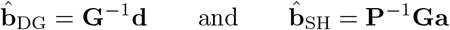

where **G** is the additive genetic variance-covariance matrix for the *n* traits and **P** is the phenotypic variance-covariance matrix. The equivalence between the two selection indices occurs when 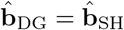, that is, when:

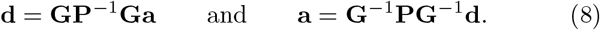

This relationship permits the expression of the desired gains in **d** with regards to the economic weights in **a** and, conversely, the definition of a desired gains vector that will yield the same profit as the Smith-Hazel Index (as demonstrated in Figure 2).

When 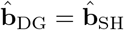, both indices are equally efficient in realising the breeding objective, whether expressed in terms of desired genetic gains or maximum profit. More specifically, both indices will now maximise the expected response to selection in the breeding values, **g**, as well as in the aggregate genotype, ℋ. Conversely, when **b**_DG_≠**b**_SH_, the Desired Gains Index will be more efficient in maximising the expected genetic gains while the Smith-Hazel Index will be more efficient in maximising profit (provided the economic weights are known without error).

#### The Desired Gains Index with BLUPs

Contemporary breeding programs generally rely on estimated breeding values (EBVs) for selecting parents for the next generation. The Desired Gains Index as proposed by Pešek and Baker (1969b), however, was designed for application with phenotypes or adjusted phenotypes. To integrate the benefits of the linear mixed model framework, we formulate a new version of the Desired Gains Index for application with BLUPs, similar to the approach which Henderson (1951) introduced for the Smith-Hazel Index.

The Desired Gains Index in Eq. 7 can be rewritten in terms of BLUPs as:

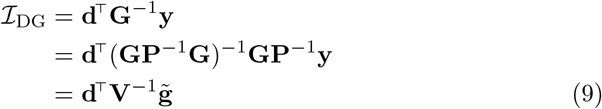

where 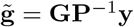 is the vector of multivariate BLUPs of the breeding values and **V** = **GP**^−1^**G** is the corresponding variance-covariance matrix, that is, 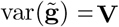. The modification of the Desired Gains Index here also maximises the expected genetic gains in proportion to the desired gains in **d**, but importantly facilitates the application of BLUPs of the breeding values instead of phenotypes. The result here is similar to the approaches of Yamada et al. (1975) and Itoh and Yamanda (1986), but note that below we explicitly show the form of index coefficients which maximise the response to selection given BLUPs of the breeding values instead of phenotypes. A brief proof of this property is provided in the following, while a general proof is derived in the Appendix.

The general selection index for an individual is given by ℐ = **b**^⊤^**y**, where **b** is the vector of selection index coefficients for the *n* traits and **y** is the vector of phenotypes. When BLUPs of the breeding values are used instead of phenotypes, the selection index can be written as 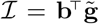, where 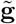 is the vector of multivariate BLUPs. The vector of expected genetic gains after one round of index selection is given by:

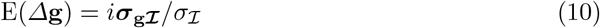

where *i* is the selection intensity, ***σ***_**g*ℐ***_ is the vector of covariances between the multivariate breeding values in **g** and the selection index 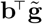, and *σ*_*ℐ*_ is the standard deviation of the selection index. These vectors are given by:

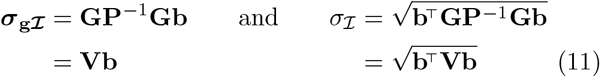

noting that 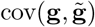 and 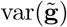 both equal **GP**^−1^**G**, that is, **V**. Since the constants *i* and *σ*_*ℐ*_ do not affect the proportionality of the index coefficients between traits, the ratio *i/σ* _ℐ_ can be set to one without loss of generality. Substituting ***σ***_**g *ℐ***_ from Eq. 11 into Eq. 10 gives:

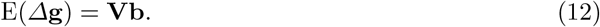

We therefore propose replacing the vector of expected genetic gains with the vector of desired genetic gains, **d**, and pre-multiplying by **V**^−1^ to obtain:

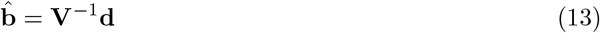

which produces the new form of the Desired Gains Index in Eq. 9 given by 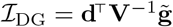.

Equivalent forms of **d** and **a** that generate identical index coefficients for the Desired Gains Index in Eq. 9 and the Smith-Hazel Index in Eq. 5 can also be obtained. The vectors of index coefficients are now given by:

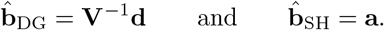

The equivalence between the two selection indices now occurs when:

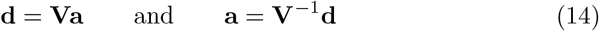

which is commensurate with the relationship in Eq. 8. This relationship again demonstrates that when 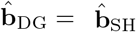, both indices are equally efficient in realising the breeding objective, whether expressed in terms of phenotypes or BLUPs.

A general form of the Desired Gains Index is presented in the Appendix, which facilitates the application of genomic estimated breeding values (GEBVs), varying amounts of information for each individual, and leverages phenotypic information from other (related) individuals, including the other selection candidates.

### 2.2 Simulations for demonstrating index selection

Stochastic simulation was used to compare four different multi-trait selection strategies for parent selection in plant breeding programs. We simulated three quantitative traits and measured genetic gain toward a predefined breeding objective across eight cycles of crossing and selection. We assumed a breeding objective that aimed to equally improve two quantitative traits while stabilising a third quantitative trait.

#### Simulation of the founder population

##### Genome simulation

Whole-genome sequences were simulated for a founder population comprising 2,000 genotypes of a hypothetical diploid line crop species. Each genome consisted of 12 homozygous chromosome pairs, with a physical length of 1 × 10^8^ base pairs (bp) and a genetic length of 100 centimorgans (cM), resulting in a total physical length of 1200 Mbp and a genetic length of 1200 cM. Chromosome sequences were generated using the Markovian coalescent simulator (Chen et al., 2008) embedded within the “runMacs2” function of AlphaSimR (Gaynor et al., 2020), with default parameters unless otherwise specified. A set of 500 biallelic quantitative trait nucleotides (QTN) and 500 single nucleotide polymorphisms (SNP) was randomly sampled along each chromosome to simulate three quantitative traits controlled by 6000 QTN and a SNP marker array with 6000 markers. Founder genotypes were formed by randomly sampling 12 chromosome pairs per genotype.

##### Simulation of genetic values and phenotypic values

Genetic values for the three quantitative traits were simulated by summing the additive genetic effects at the 6000 QTN. The additive genetic effects were randomly sampled from a multivariate standard normal distribution and scaled to an additive genetic variance of 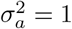 in the founder population for all three traits. The genetic correlation matrix for the three traits was set as:

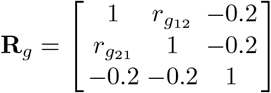

where *r*_*g*12_ = *r*_*g*21_ was set as 0.4, 0, or -0.4.

Phenotypes were generated by adding random errors to the genetic values. The random errors were sampled from a standard normal distribution and scaled to a trait-specific error variance 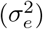 to obtain heritabilities of 0.2, 0.7, and 0.4 in the founder population for the three traits, respectively. The heritabilities were calculated as the ratio of additive genetic to total phenotypic variance given by 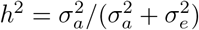.

#### Simulation of multi-trait testing strategies

We conducted simulations across eight cycles of crossing and selection to compare four different multi-trait selection strategies for parent selection. Our breeding objective aimed to equally improve traits 1 and 2 while maintaining trait 3 at a stable value of 0. The selection strategies encompassed i) a Desired Gains Index, ii) a Smith-Hazel Index, iii) independent culling, and iv) selection of extreme genotypes. Each cycle involved selecting 30 new parents, which were then crossed to produce 100 bi-parental F1 genotypes. From each F1, 20 homozygous full-sibs were derived to replicate a single-seed descent breeding scheme for genotype fixation through repeated selfing. Genetic gain was assessed as the mean genetic value of the breeding population over the eight cycles of crossing and selection. Each evaluation comprised 100 simulation replicates. Additionally, we measured genetic variance and parent selection accuracy (Supplementary S1-S6), with selection accuracy measured as the Pearson correlation between the true genetic values and the phenotypes or the genomic estimated breeding values (GEBVs) of the selection candidates.

##### Parent selection using a Desired Gains Index

Parent selection using the Desired Gains Index aimed to achieve an improvement ratio of 1:1:0 for traits 1, 2, and 3, respectively. To accomplish this, the desired gains vector, **d**, was recalculated at each cycle with reference to the realised gains in the population. The top 30 parents with the highest index were randomly crossed to generate the F1 genotypes. To demonstrate the superiority of a Desired Gains Index with BLUPs over (adjusted) phenotypes, we compared selection based on phenotypes to selection using GEBVs obtained from a multivariate genomic prediction model. For simplicity, we used the true additive genetic variance-covariance matrix, **G**, to compute the weights with phenotypes, potentially leading to a slight inflation in observed gains and accuracies. Additionally, parent selection using a Desired Gains Index with true genetic values, **g**, and the true additive genetic variance-covariance matrix, was included as a control scenario.

##### Parent selection using the Smith-Hazel Index

Parent selection using the Smith-Hazel index employed economic weights of 1:1:0 for traits 1, 2, and 3, respectively. Our objective was to demonstrate the effect of a vector **a** = (1, 1, 0)^⊤^ with economic weights equal to the desired gains improvement ratio. The top 30 parents with the highest index were randomly crossed to generate the F1 genotypes. All selections were based on GEBVs obtained from a multivariate genomic prediction model.

##### Parent selection using a Desired Gains Index with d = Va

To demonstrate the equivalence between the Desired Gains Index and Smith-Hazel Index, parent selection using the Desired Gains Index was also implemented with a vector of desired gains derived from the vector of economic values. The vector of desired gains was calculated as **d** = **Va**, where **V** is the variance-covariance matrix of the GEBVs and **a** = (1, 1, 0)^⊤^. The top 30 parents with the highest index were randomly crossed to generate the F1 genotypes. All selections were based on GEBVs obtained from a multivariate genomic prediction model.

##### Parent selection using independent culling

Parent selection using independent culling followed a step-wise approach. Initially, our objective was stabilising trait 3 by selecting 30% of the population with values close to zero. Next, we identified the top 10% genotypes for trait 1 from the remaining population, followed by selecting 30 parents based on superior trait 2 values. Parents were then randomly crossed to generate the F1. To demonstrate the influence of selection order on genetic gain, we repeated this process with trait 2 selection preceding trait 1. All selections were based on GEBVs obtained from a multivariate genomic prediction model.

##### Selection of extreme genotypes as parents

Selection of extreme genotypes as parents followed a step-wise approach. Initially, our objective was stabilising trait 3 by selecting 30% of the population with values close to zero. Next, we selected and removed the top 15 genotypes for trait 1 from the remaining population. Subsequently, we selected the 15 genotypes with the highest values for trait 2. From all possible 225 crosses between the two sets of 15 parents, 100 crosses were randomly selected to generate the F1. All selections were based on GEBVs obtained from a multivariate genomic prediction model.

## 3 Results

### 3.1 Parent selection using a Desired Gains Index with phenotypes and GEBVs

The Desired Gains Index (DGI) with GEBVs exhibited superior performance compared to the DGI with phenotypes. This is shown in the first two columns of Figure 1, which plots genetic gain against time for the three simulated traits. Both methods equally enhanced traits 1 and 2 while maintaining trait 3 close to zero. However, genetic gains for traits 1 and 2 were substantially higher when using the DGI with GEBVs (Table 1). For example, at a genetic correlation of 0.4 between trait 1 and trait 2, the DGI with GEBVs demonstrated 20% more genetic gain for traits 1 and 2 than the DGI with phenotypes.

**Table 1.**
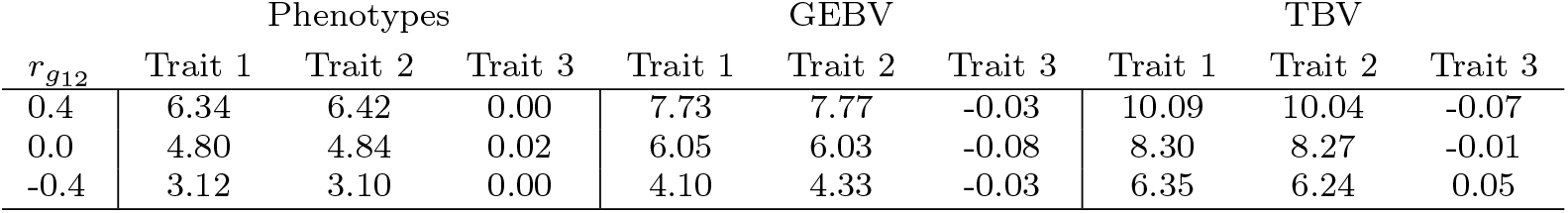
Mean realised genetic gain after eight cycles of crossing and selection using a Desired Gains Index with phenotypes, genomic estimated breeding values (GEBV), and true breeding values (TBV). Genetic gain is presented for pairwise genetic correlations (*r*_*g*12_) between trait 1 and trait 2 of 0.4, 0, and -0.4, respectively.

**Fig. 1.**
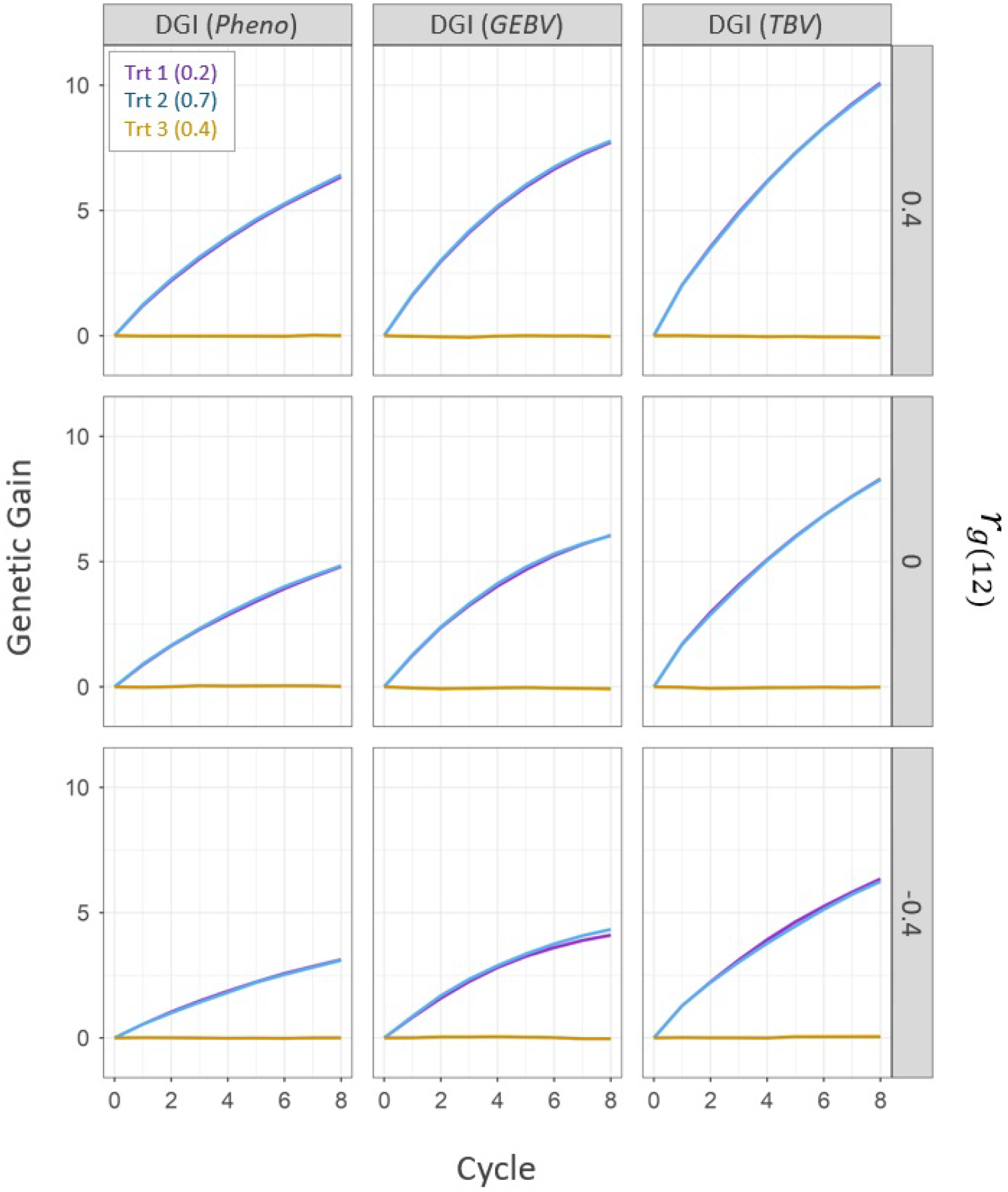
Genetic gain for three simulated quantitative traits over eight cycles of crossing and parent selection using a Desired Gains Index (DGI) with: i) phenotypes (pheno), ii) genomic estimated breeding values (GEBV), and iii) simulated true breeding values (TBV). In each panel, genetic gain is plotted as the change in the mean additive genetic value over time. Trait 1 is represented by the purple line (Trt 1), trait 2 by the blue line (Trt 2), and trait 3 by the yellow line (Trt 3). Numbers in parentheses indicate the simulated trait heritabilities (*h*^2^) in the founder population. Each of the three Desired Gains Index selection scenarios (columns) was evaluated under a pairwise genetic correlation (**r_g12_**) between trait 1 and trait 2 of 0.4, 0, and -0.4 (rows), respectively. The pairwise genetic correlations between trait 1 and trait 3, and between trait 2 and trait 3, were set to -0.2. The breeding objective was to achieve an improvement ratio of 1:1:0 for traits 1, 2, and 3, respectively.

Figure 1 also shows a decrease in genetic gain for traits 1 and 2 as the genetic correlation between the two traits decreases. For example, at a genetic correlation of -0.4, the DGI with GEBVs exhibited almost 50% less gain for trait 1 and trait 2, respectively, compared to a genetic correlation of 0.4. However, lower genetic correlations resulted in an increased relative advantage of the DGI with GEBVs over the DGI with phenotypes (Table 1).

As expected, the control scenario with parent selection using a DGI with true breeding values (TBV) showed the best performance, indicating the upper efficiency limit when h2 = 1.

### 3.2 Parent selection using a Smith-Hazel Index

The application of the Smith-Hazel index (SHI) with the economic weights vector **a** = (1, 1, 0)^⊤^ did not yield an effective strategy for parent selection to meet our breeding objective. This is shown in the second column of Figure 2, which plots genetic gain against time for the three simulated traits. Across all three genetic correlations, trait 1 exhibited greater genetic gain than trait 2, while trait 3 showed negative genetic gain. This outcome is expected, as the 1:1:0 economic weight ratio employed with the SHI has different implications for selection compared to the desired gains improvement ratio of 1:1:0, as discussed further below.

**Fig. 2.**
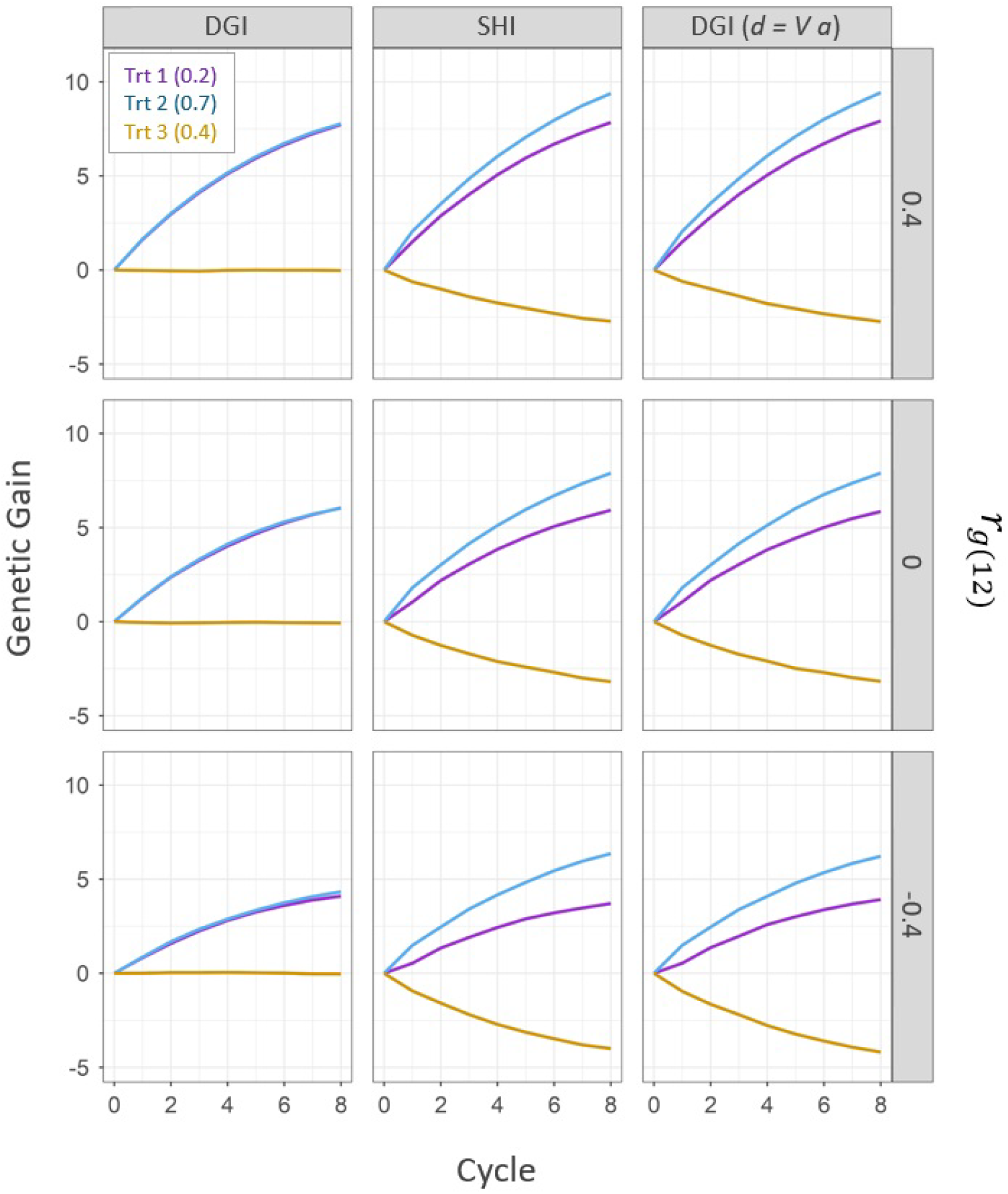
Genetic gain for the three simulated quantitative traits over eight cycles of crossing and parent selection using a Desired Gains Index (DGI) and a Smith-Hazel Index (SHI) with genomic estimated breeding values. In each panel, genetic gain is plotted as the change in the mean additive genetic value over time. Trait 1 is represented by the purple line (Trt 1), trait 2 by the blue line (Trt 2), and trait 3 by the yellow line (Trt 3). Numbers in parentheses indicate the simulated trait heritabilities (*h*^2^) in the founder population. For the Desired Gains Index (column 1), the targeted improvement ratio was set to 1:1:0 for traits 1, 2, and 3, respectively. For the Smith-Hazel Index (column 2), the economic weights in vector **a** were set to 1:1:0 for traits 1, 2, and 3, respectively. The comparison of the two scenarios illustrates that improving genetic gain and improving economic value are two fundamentally different breeding objectives. However, to demonstrate the equivalence of the Desired Gains Index and the Smith-Hazel Index, a Desired Gains Index with the improvement ratio derived from the vector of economic values **a** (column 3) was simulated (see Eq. 14). Each of the selection scenarios (columns) was evaluated under a pairwise genetic correlation (**r_g12_**) between trait 1 and trait 2 of 0.4, 0, and -0.4 (rows), respectively. The pairwise genetic correlations between trait 1 and trait 3, and between trait 2 and trait 3, were set to -0.2.

Moreover, as shown in the third column of Figure 2, genetic gains similar to the SHI can be achieved using a DGI, given the vector of desired gains is computed as **d** = **Va**, where **V** represents the covariance matrix of the GEBVs.

### 3.3 Parent selection using independent culling and selection of extreme genotypes

Independent culling and selection of extreme genotypes exhibited significantly poorer performance in achieving our breeding objective than the DGI. This is shown in Figure 3, which plots genetic gain against time for the three simulated traits. Our breeding objective aimed to equally enhance trait 1 and trait 2 while maintaining trait 3 at a stable value of zero. Only the DGI consistently achieved this at all three genetic correlations. Independent culling and selection of extreme genotypes managed to maintain trait 3 close to zero. However, the two parent selection strategies demonstrated notably different genetic gain for traits 1 and trait 2. While selection of extreme genotypes showed a somewhat balanced improvement of both traits, genetic gains were substantially lower than those achieved with the DGI. Independent culling, on the other hand, failed to achieve a balanced improvement of both traits. Specifically, when the genetic correlations decreased, the difference in gain between trait 1 and trait 2 became strongly pronounced.

**Fig. 3.**
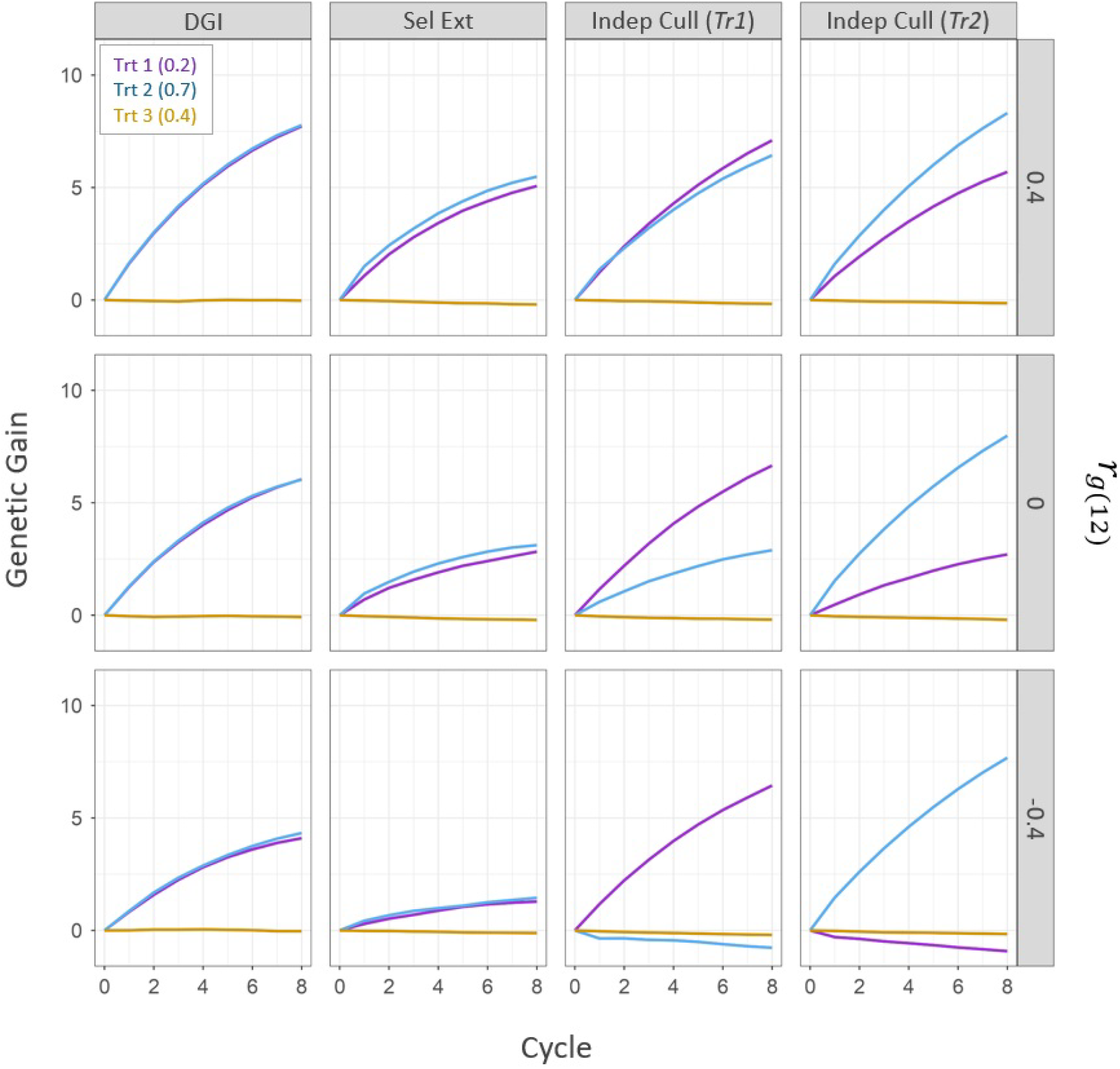
Genetic gain for the three simulated quantitative traits over eight cycles of crossing and parent selection using a Desired Gains Index (DGI), selection of extreme genotypes (Sel Ext), and independent culling (Indep Cull) with genomic estimated breeding values. In each panel, genetic gain is plotted as the change in the mean additive genetic value over time. Trait 1 is represented by the purple line (Trt 1), trait 2 by the blue line (Trt 2), and trait 3 by the yellow line (Trt 3). Numbers in parentheses indicate the simulated trait heritabilities (*h*^2^) in the founder population. The targeted improvement ratio was 1:1:0 for traits 1, 2, and 3, respectively. For selection of extremes (column 2), initially, trait 3 was stabilised by selecting 30% of the population with values close to zero. Then, the best 15 genotypes were selected for traits 1 and trait 2, respectively (without overlap), and from all possible 225 crosses, 100 were randomly selected to generate the F1. Parent selection using independent culling (columns 3 4) followed a step-wise approach. Initially, trait 3 was stabilised by selecting 30% of the population with values close to zero. Then, the top 10% genotypes for trait 1 were identified from the remaining population, followed by selecting 30 parents based on superior trait 2 values (*Tr1*). Parents were then randomly crossed to generate the F1. To demonstrate the influence of selection order on genetic gain, this process was repeated with trait 2 selection preceding trait 1 (*Tr2*). Each of the selection scenarios (columns) was evaluated under a pairwise genetic correlation (**r_g12_**) between trait 1 and trait 2 of 0.4, 0, and -0.4 (rows), respectively. The pairwise genetic correlations between trait 1 and trait 3, and between trait 2 and trait 3, were set to -0.2.

Figure 3 also shows that when independent culling was used, the order of trait selection significantly influenced the genetic gain of trait 1 and trait 2. Notably, the trait chosen first consistently exhibited higher genetic gain.

## 4 Discussion

### 4.1 The Desired Gains Index with BLUPs is better than with phenotypes

The Desired Gains Index with BLUPs, or more specifically GEBVs, resulted in more genetic gain for the two improvement traits than the index with phenotypes. The methods developed in this paper show how the BLUP framework can be leveraged for the Desired Gains Index, in a similar manner to what Henderson (1951) showed for the Smith-Hazel Index. There are three distinct advantages of this approach. Firstly, the BLUPs are commonly obtained from a linear mixed model which can accommodate many routine features of plant breeding data, such as unbalanced and incomplete experimental designs and complex genetic and non-genetic variance structures. Secondly, they can utilise varying amounts of information from other related individuals to improve the accuracy of the selection index, e.g., by utilising relevant phenotypic data across multiple years and stages. Finally, they enable seamless integration with genomic selection, allowing the utilisation of GEBVs, as demonstrated here. The Desired Gains Index with BLUPs developed in this paper therefore provides opportunities to leverage many different data sources and many different modern plant breeding tools.

### 4.2 The Smith-Hazel index maximises economic value, not genetic gain

The Smith-Hazel index aims to maximise the economic value of the progeny population, not genetic gain. Although this is widely acknowledged, its crucial to recognise the distinction between a ratio of economic weights and a desired gains improvement ratio.

Unlike the Desired Gains Index, which prioritises approximation of the desired improvement ratio in the next generation, the Smith-Hazel index focuses on improving traits with high economic values. Economic gain, however, does not necessarily translate into genetic gain, and traits with low economic value may be sacrificed to improve those with higher economic value. In fact, poorly chosen economic weights can lead to negative genetic gains in traits intended to exhibit positive gains. For example, the Smith-Hazel index might prioritise maximizing gain in a trait with high economic weight, such as yield, even if it means sacrificing genetic gain in a trait with a potentially lower but positive economic weight, such as oil or protein content in the seed. Likewise, an economic value of zero (or no assigned value) does not facilitate trait stabilisation but indicates that changes in the trait value hold no significance for the final product. To balance profit maximisation in some traits with stabilising other traits, more complex modifications of the Smith-Hazel Index are required (see Brascamp, 1984).

Furthermore, trait heritability has an impact on which traits are improved. For example, suppose two traits have similar economic values but differ in heritabilities. Given the trait with higher heritability’s greater potential for genetic gain, the Smith-Hazel index will allocate greater importance to it. It should be noted, however, that neither the Desired Gains Index nor the Smith-Hazel Index can per se be considered better since they pursue fundamentally different breeding objectives. Both approaches, however, maximise the expected genetic gains in proportion to the breeding objective and can, therefore, be considered optimal.

### 4.3 Independent culling and selection of extreme genotypes are inefficient methods for multi-trait parent selection

Independent culling and selection of extreme genotypes are generally considered less efficient for multi-trait parent selection compared to index selection (e.g., Hazel and Lush, 1942; Young, 1961; Pešek and Baker, 1969a; Batista et al., 2021). Both strategies, however, prove valuable at different stages of plant breeding when applied appropriately.

Independent culling is the practice of setting minimum thresholds (culling levels) for various traits and removing individuals that fail to meet these thresholds for all traits of interest. The thresholds are typically determined based on farmer needs, market demands, or in comparison with benchmark agronomic checks. Independent culling is particularly useful during product development stages, after parental selection, to identify candidate varieties that meet all minimum requirements for product registration and release. In contrast, selection indices like the Desired Gains Index and the Smith-Hazel Index are unsuitable at this stage, as they are designed for population improvement rather than meeting minimum thresholds.

In practice, parent selection often employs a cost-effective combination of independent culling and index selection. Quantitative traits with high heritability, such as plant height or flowering time, can be assessed early, allowing genotypes outside the desired phenotypic range to be discarded. Similarly, independent culling may be employed for qualitative traits, such as major resistance genes, through marker-assisted selection. Following this initial culling process, index selection is applied to the remaining subset. While this approach may not optimize genetic gains in all traits to their fullest potential, the logistical and cost constraints inherent in plant breeding programs necessitate a somewhat pragmatic approach that balances costs and gains effectively. Figure 4 illustrates how independent culling and index selection can be efficiently integrated into a line breeding program, loosely based on the structure of the CGIAR’s wheat breeding programs.

**Fig. 4.**
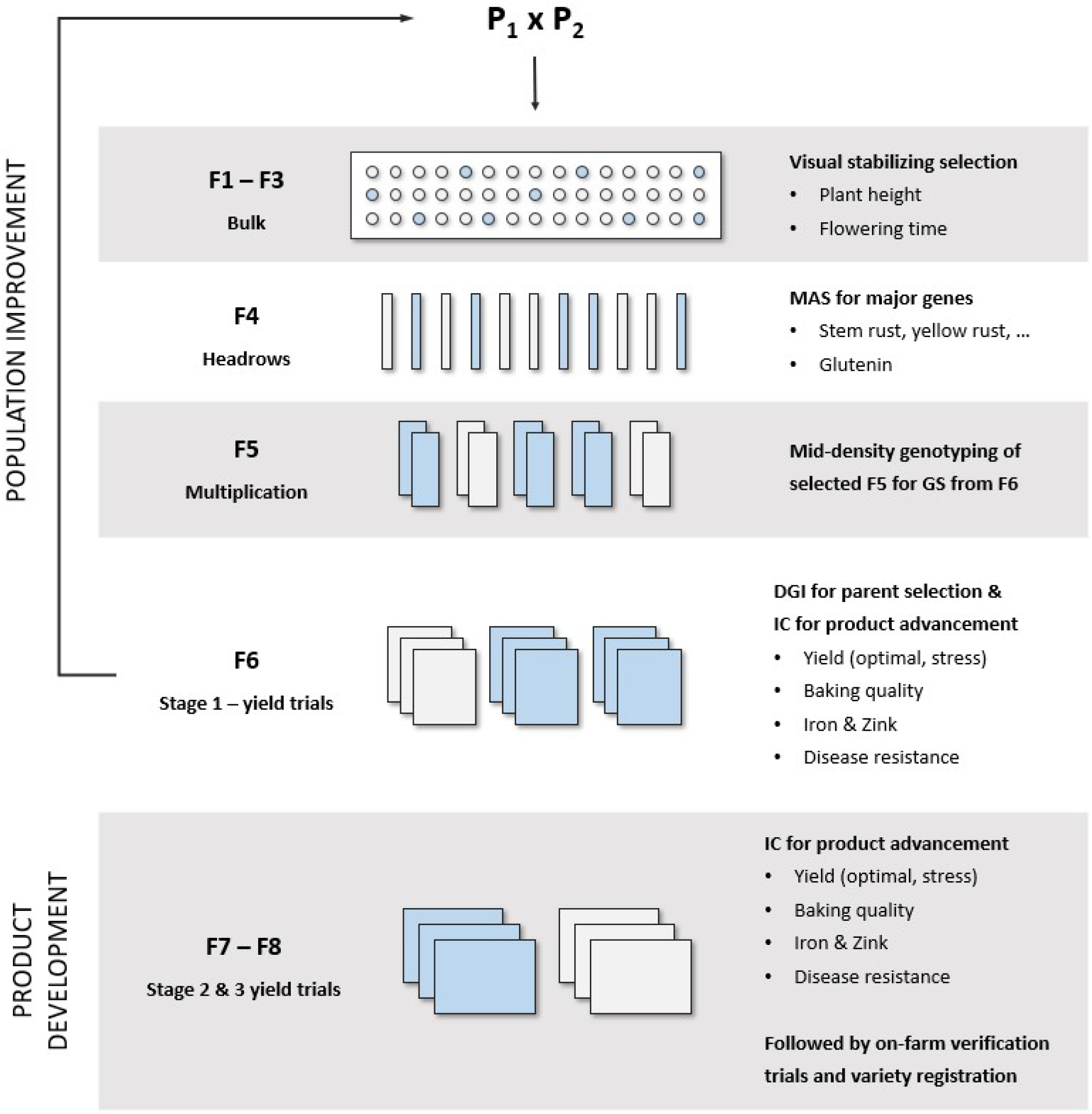
Schematic representation of a line breeding program efficiently combining Desired Gains Index (DGI) selection for population improvement and independent culling (IC) for product development. The structure of the presented line breeding program is loosely based on the structure of the CGIAR’s wheat breeding programs.

Extreme genotypes are often used to combine desirable traits from two distinct parents in the progeny generation. While this approach may seem intuitive, our simulations demonstrate its limitations, particularly with quantitative traits. Efficiently improving two traits through crossing parents with strong performance in one trait each is exceptionally difficult, particularly when genetic correlations between traits are low or negative. This challenge is intensified when dealing with more than two or three traits characterised by diverse pairwise genetic correlations. Despite its limitations with quantitative traits, crossing extreme genotypes is a valid strategy for introgressing important genes into elite backgrounds lacking those genes while maintaining high performance in other traits. However, this typically involves multiple cycles of backcrossing to restore as much of the elite genetic background as possible around the introgressed gene or genes. Thus, it falls within the domain of “trait development and deployment” (or “pre-breeding”) rather than population improvement.

### 4.4 Understanding the genetic correlations between traits can enable the improvement of the breeding objective

The Desired Gains Index is an optimal solution for parent selection in plant breeding programs when the breeding objectives are defined in terms of desired response to selection for multiple traits. Using a desired gains index ensures that population improvement progresses at the desired improvement ratio over time. However, the rate of genetic gain for each trait, and thus the time until the breeding objective is achieved, is determined by the heritabilities of and pairwise genetic correlations between the traits. Unfavourable genetic correlations between two or more traits can impede progress, especially if the objective is to enhance those traits at an equal rate. Hence, thoroughly investigating the relationship between genetic correlations and the expected genetic gain space across multiple traits often proves advantageous. For instance, sacrificing a negligible amount of potential gain in one trait could lead to significantly greater gain in one or several other traits. Drawing on selection theory, software tools such as DESIRE (Kinghorn, 2013) empower breeders to fine-tune their expected genetic gains and potentially optimise their breeding objectives.

### 4.5 Simulating fully pleiotropic traits provides a conservative estimate of gains across multiple traits

A limitation of AlphaSimR is its requirement for multiple traits with a predefined genetic correlation to be completely pleiotropic. This implies that the genetic correlation between traits is fully determined by the simulated effects at all quantitative trait loci, representing a scenario where genetic correlations between traits cannot be broken down.

Genetic correlations can arise from two factors: linkage disequilibrium and pleiotropy (Hill, 2013). Genetic correlations are dynamic and subject to change over time due to selection and recombination. While genetic correlations stemming from linkage disequilibrium can be broken down by repeated cycles of meiosis (Bernardo, 2010), they may become increasingly negative over time due to selection, known as the Bulmer effect (Bulmer, 1971; Batista et al., 2021). Low or negative genetic correlations between traits diminish the efficacy of selection on multiple traits, as improving one trait may restrict gains in others. In our simulations, this limitation in gain was fixed and irremovable, whereas, in reality, various evolutionary forces, such as recombination and selection, may facilitate multi-trait selection over time when traits are negatively correlated.

### 4.6 Concluding remarks

The Desired Gains Index is an optimal solution for parent selection in plant breeding programs where the breeding objective is defined in terms of desired response to selection. To facilitate its implementation within the framework of modern statistical analysis tools, we have provided a derivation of the Desired Gains Index using Best Linear Unbiased Predictors (BLUPs). This also provides the foundation for the integration of the Desired Gains Index with rapid recycling genomic selection strategies, such as the two-part breeding strategy (e.g., Gaynor et al., 2017; Powell et al., 2020; Werner et al., 2023). Additionally, the Desired Gains Index can seamlessly complement other strategies for optimizing parent selection, such as optimum contribution selection to harmonize long-term genetic gain and genetic variance (Woolliams et al., 2015; Gorjanc et al., 2018).

## Statements & Declarations

### Author Contributions

CW and KG conceived the study. CW and DT developed the methodology, carried out the simulations, and wrote the manuscript. DT derived the general forms in the Appendix.

### Data Availability

The code and datasets simulated for the current study are available from the corresponding author on request.

## Appendix A General forms of the selection indices

This appendix derives general forms of the Smith-Hazel Index and Desired Gains Index for application with Best Linear Unbiased Predictors (BLUPs). The objective here is to provide general forms which handle genomic estimated breeding values (GEBVs) and highlight how the required components are readily available from the fit of a multivariate linear mixed model. Simplifications of the methods developed here are presented in the Introduction and Materials and methods.

### A.1 Preliminaries

Assume that phenotypes are available on *n* traits for *v* individuals, that is, the selection candidates. The application to other (related) individuals with or without phenotypic information is straightforward. Let the *η*-vector of phenotypic data for all individuals be given by 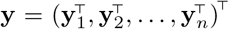, where **y**_*j*_ is the *η*_*j*_-vector for trait *j*. The multivariate linear mixed model for **y** can be written as:

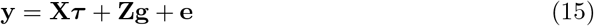

where **τ** is a *p*-vector of fixed effects, including the trait means, with design matrix **X** (assumed to have full column rank), **g** is the *nv*-vector of random multivariate breeding values with design matrix **Z** and **e** is the *η*-vector of random non-genetic effects and residuals. It is assumed that:

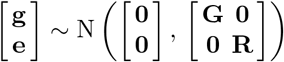

where **G** is the *nv* × *nv* genetic variance-covariance matrix and **R** is the *η* × *η* residual variance-covariance matrix. Both **G** and **R** are assumed to be completely general, but note that **G** captures the genetic covariances between traits as well as the relationships between individuals, for example, through genomic or pedigree relationship information.

Let **g**_*i*_ denote the *n*-vector of multivariate breeding values for individual *i*. It then follows that:

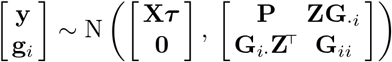

where **P** = **ZGZ**^⊤^ + **R** is the *η* × *η* phenotypic variance-covariance matrix (assumed to be non-singular), **G**_*ii*_ is a *n* × *n* matrix containing the genetic variances and co-variances of individual *i* and **G**_*i*·_ is a *n* × *nv* matrix containing the genetic variances and covariances of individual *i* and all other individuals, that is, var(**g**_*i*_, **g**) = **G**_*i*·_.

Standard linear mixed model results give:

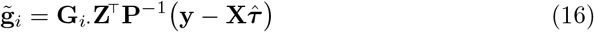

where 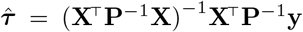 is the Best Linear Unbiased Estimator (BLUE) of the fixed effects. The prediction error variance-covariance matrix of 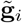 is obtained as:

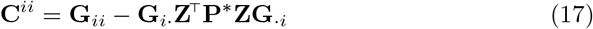

where **P**^*^ = **P**^−1^ − **P**^−1^**X**(**X**^⊤^**P**^−1^**X**)^−1^**X**^⊤^**P**^−1^, such that 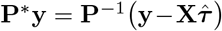. The matrix **C** is the part of the inverse coefficient matrix corresponding to 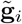, which is routinely provided by linear mixed model software. The variance-covariance matrix of 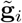 is then obtained as **V**_*ii*_ = **G**_*ii*_ − **C**^*ii*^, that is, **V**_*ii*_ = **G**_*i*·_**Z**^⊤^**P**^*^**ZG**_·*i*_.

### A.2 The Smith-Hazel Index

Let the aggregate genotype for individual *i* be given by ℋ_*i*_ = **a**^⊤^**g**_*i*_, where **a** is the *n*-vector of economic weights for the traits and **g**_*i*_ is the *n*-vector of multi-variate breeding values. The general selection index is calculated as:

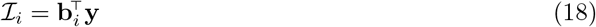

where **b**_*i*_ is the *η* vector of selection index coefficients and **y** is the *η*-vector of phenotypes. It is important to note that the general form of ℐ_*i*_ developed here allows for all (related) individuals with phenotypic data to contribute to the selection index of individual *i*, including the other selection candidates. This allows for information sharing across individuals as well as traits.

The expected genetic gain in the aggregate genotype after one cycle of index selection is given by:

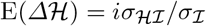

where *i* is the selection intensity, *σ* _ℋ ℐ_ is the covariance between ℋ _*i*_ and ℐ_*i*_ given by **a**^⊤^**G**_*i*·_**Z**^⊤^**b**_*i*_ and *σ*_I_ is the standard deviation of ℐ _*i*_ given by the square-root of 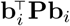. Following Henderson (1963), E(*Δ*ℋ) is maximised subject to the constraint E(ℐ) = 0, which produces the set of index equations:

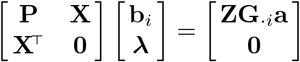

where ***λ*** is a *p*-vector of Lagrange multipliers. Solving for **b**_*i*_ gives:

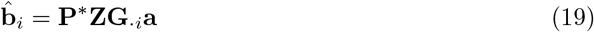

where **P**^*^ = **P**^−1^ **P**^−1^**X**(**X**^⊤^**P**^−1^**X**)^−1^**X**^⊤^**P**^−1^ from Eq. 17. Substituting 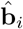 into Eq. 18 produces a general form of the Smith-Hazel Index given by:

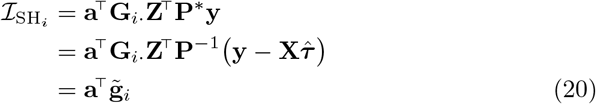

where 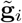 is the *n*-vector of multivariate BLUPs of the breeding values from Eq. 16. The Smith-Hazel Index therefore maximises the expected response to index selection in ℋ_*i*_ in relation to the economic weights in **a**. Note, however, this is only true when the economic weights are known without error.

The Smith-Hazel Index in Eq. 20 is an extension of the methods in Henderson (1963) for use with GEBVs, or more generally BLUPs involving relationships between the selection candidates. In practice, the selection index is formed simultaneously for all selection candidates as 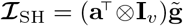, where 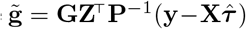 is the *nv*-vector of multivariate BLUPs for all individuals.

### A.3 The Desired Gains Index

Recall the vector of multivariate breeding values for individual *i* is given by **g**_*i*_ and the general selection index is calculated as 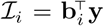. The expected genetic gains in the vector of breeding values after one cycle of index selection is given by:

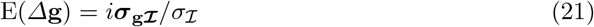

where *i* is the selection intensity, ***σ***_**g*ℐ***_ is the *n*-vector of covariances between **g**_*i*_ and **ℐ** _*i*_ given by **G**_*i*·_**Z**^⊤^**b**_*i*_ and *σ***ℐ** is the standard deviation of **ℐ**_*i*_ given by the square-root of 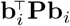. Pešek and Baker (1969b) and Yamada et al. (1975) proposed replacing the vector of expected genetic gains in Eq. 21 with a vector of desired genetic gains chosen here to satisfy E(*Δ***g**) = *i***d***/σ*_*I*_, that is, cov(**g**, ℐ) = **d**.

Extending Itoh and Yamanda (1986), E(*Δ***g**) is maximised subject to the constraints cov(**g**, ℐ) = **d** and E(ℐ) = 0, which produces the set of index equations:

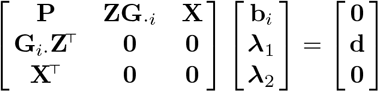

where ***λ***_1_ is a *n*-vector of Lagrange multipliers and ***λ***_2_ is a *p*-vector of Lagrange multipliers. Solving for **b**_*i*_ gives:

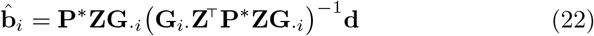

where **P**^*^ = **P**^−1^ **P**^−1^**X**(**X**^⊤^**P**^−1^**X**)^−1^**X**^⊤^**P**^−1^ from Eq. 17. Substituting 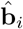 into Eq. 18 produces a general form of the Desired Gains Index given by:

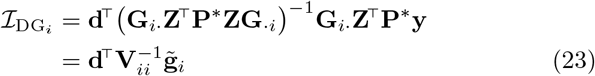

where 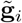 is the *n*-vector of multivariate BLUPs of the breeding values from Eq. 16 and **V**_*ii*_ is the corresponding variance-covariance matrix. The Desired Gains Index therefore maximises the expected response to index selection in **g**_*i*_ in proportion to the desired genetic gains in **d**.

The Desired Gains Index in Eq. 23 is an extension of the methods in Yamada et al. (1975) and Itoh and Yamanda (1986) for use with GEBVs, or more generally, BLUPs involving relationships between the selection candidates. The selection index can be formed simultaneously for all selection candidates as 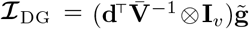, where 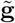 is the *nv*-vector of multivariate BLUPs for all individuals and 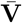 is the *n* × *n* marginal variance-covariance matrix given by 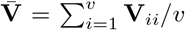, assumed to be non-singular. We have found that this provides a good approximation to the full selection index for balanced data (see the Supplementary Material).

### A.4 General relationship between the Desired Gains Index and the Smith-Hazel Index

The equivalence between the Desired Gains Index and the Smith-Hazel Index occurs when the index coefficients in Eqs. 19 and 22 are equal, that is when:

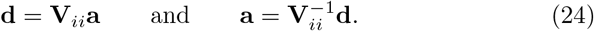

This relationship permits the expression of the desired gains in **d** with regards to the economic weights in **a**, and vice versa, such that both indices are equally efficient at improving genetic gain and maximising profit. It is important to note that the equivalence requires a separate expression for each individual. When the Desired gains index is approximated using the marginal variance matrix, 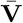, the equivalence between the selection indices occurs when 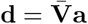 and 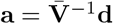 for all individuals.

## Supplementary material

**Fig. S1.**
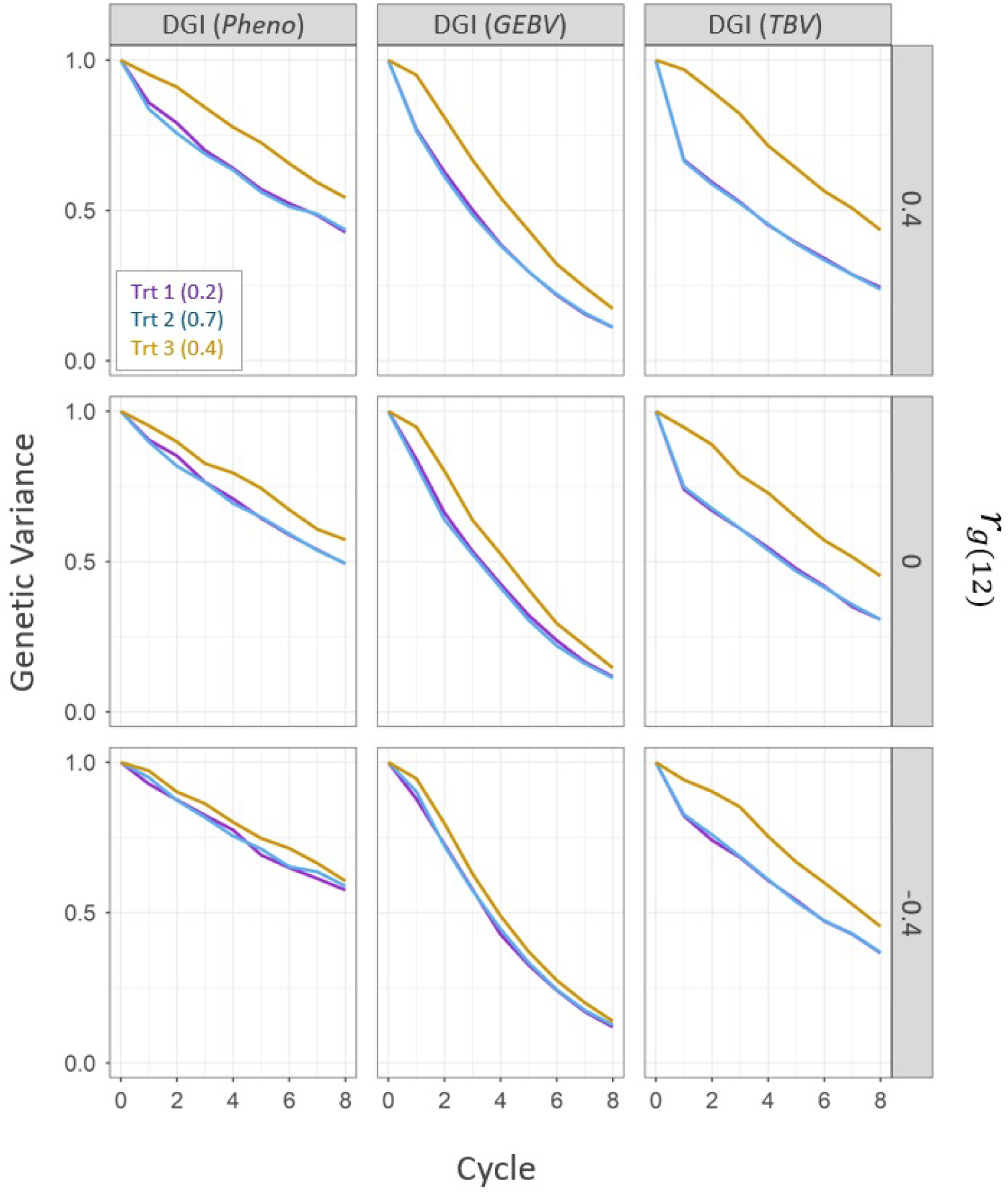
Genetic variance for three simulated quantitative traits over eight cycles of crossing and parent selection using a Desired Gains Index (DGI) with: i) phenotypes (pheno), ii) genomic estimated breeding values (GEBV), and iii) simulated true breeding values (TBV). In each panel, genetic variance is plotted as the change in the mean additive genetic variance over time. Trait 1 is represented by the purple line (Trt 1), trait 2 by the blue line (Trt 2), and trait 3 by the yellow line (Trt 3). Numbers in parentheses indicate the simulated trait heritabilities (*h*^2^) in the founder population. Each of the three Desired Gains Index selection scenarios (columns) was evaluated under a pairwise genetic correlation (**r_g12_**) between trait 1 and trait 2 of 0.4, 0, and -0.4 (rows), respectively. The pairwise genetic correlations between trait 1 and trait 3, and between trait 2 and trait 3, were set to -0.2. The breeding objective was to achieve an improvement ratio of 1:1:0 for traits 1, 2, and 3, respectively.

**Fig. S2.**
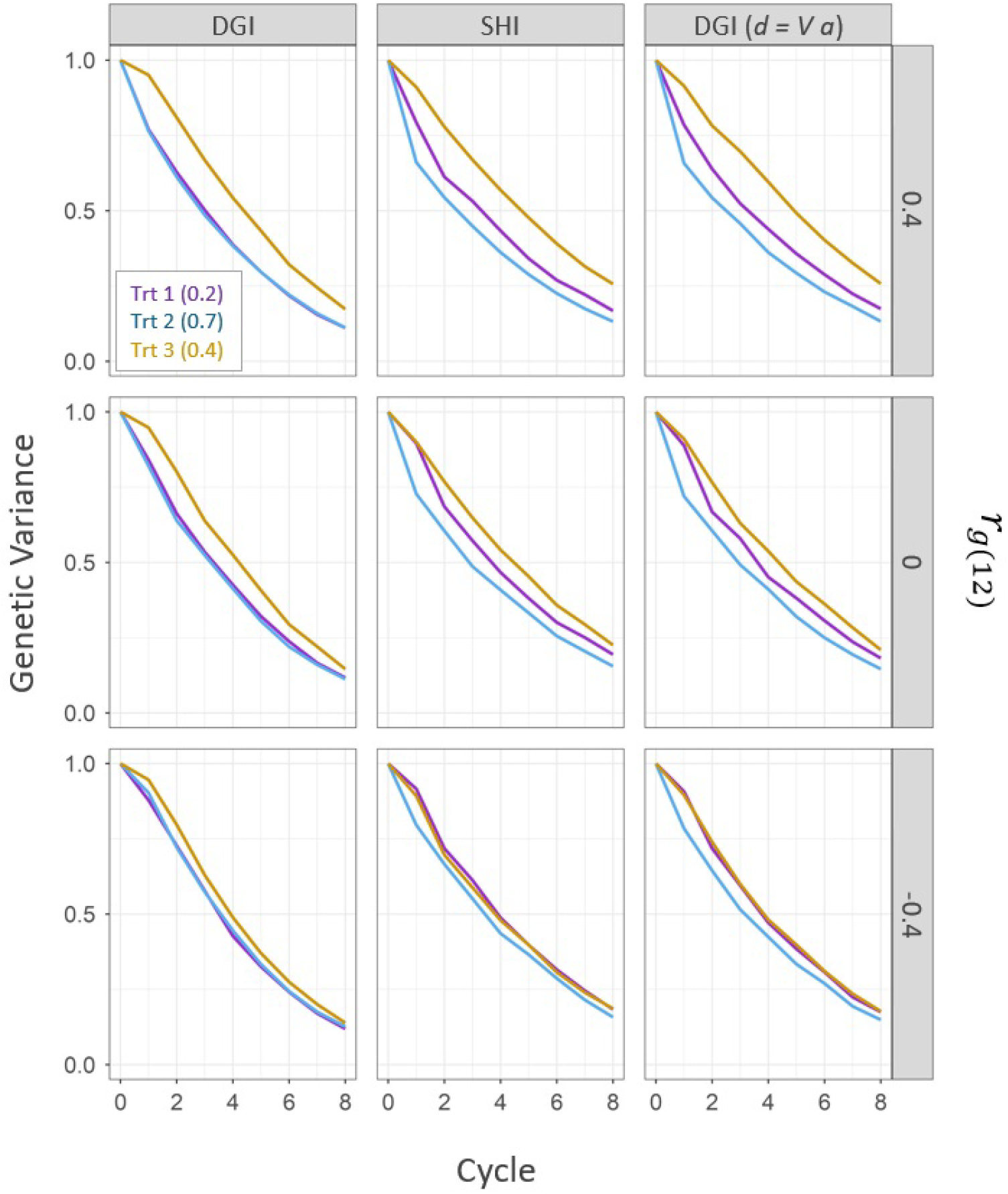
Genetic variance for the three simulated quantitative traits over eight cycles of crossing and parent selection using a Desired Gains Index (DGI) and a Smith-Hazel Index (SHI) with genomic estimated breeding values. In each panel, genetic variance is plotted as the change in the mean additive genetic variance over time. Trait 1 is represented by the purple line (Trt 1), trait 2 by the blue line (Trt 2), and trait 3 by the yellow line (Trt 3). Numbers in parentheses indicate the simulated trait heritabilities (*h*^2^) in the founder population. For the Desired Gains Index (column 1), the targeted improvement ratio was set to 1:1:0 for traits 1, 2, and 3, respectively. For the Smith-Hazel Index (column 2), the economic weights in vector **a** were set to 1:1:0 for traits 1, 2, and 3, respectively. Column 3 shows a Desired Gains Index with the improvement ratio derived from the vector of economic values **a** (see Eq. 14). Each of the selection scenarios (columns) was evaluated under a pairwise genetic correlation (**r_g12_**) between trait 1 and trait 2 of 0.4, 0, and -0.4 (rows), respectively. The pairwise genetic correlations between trait 1 and trait 3, and between trait 2 and trait 3, were set to -0.2.

**Fig. S3.**
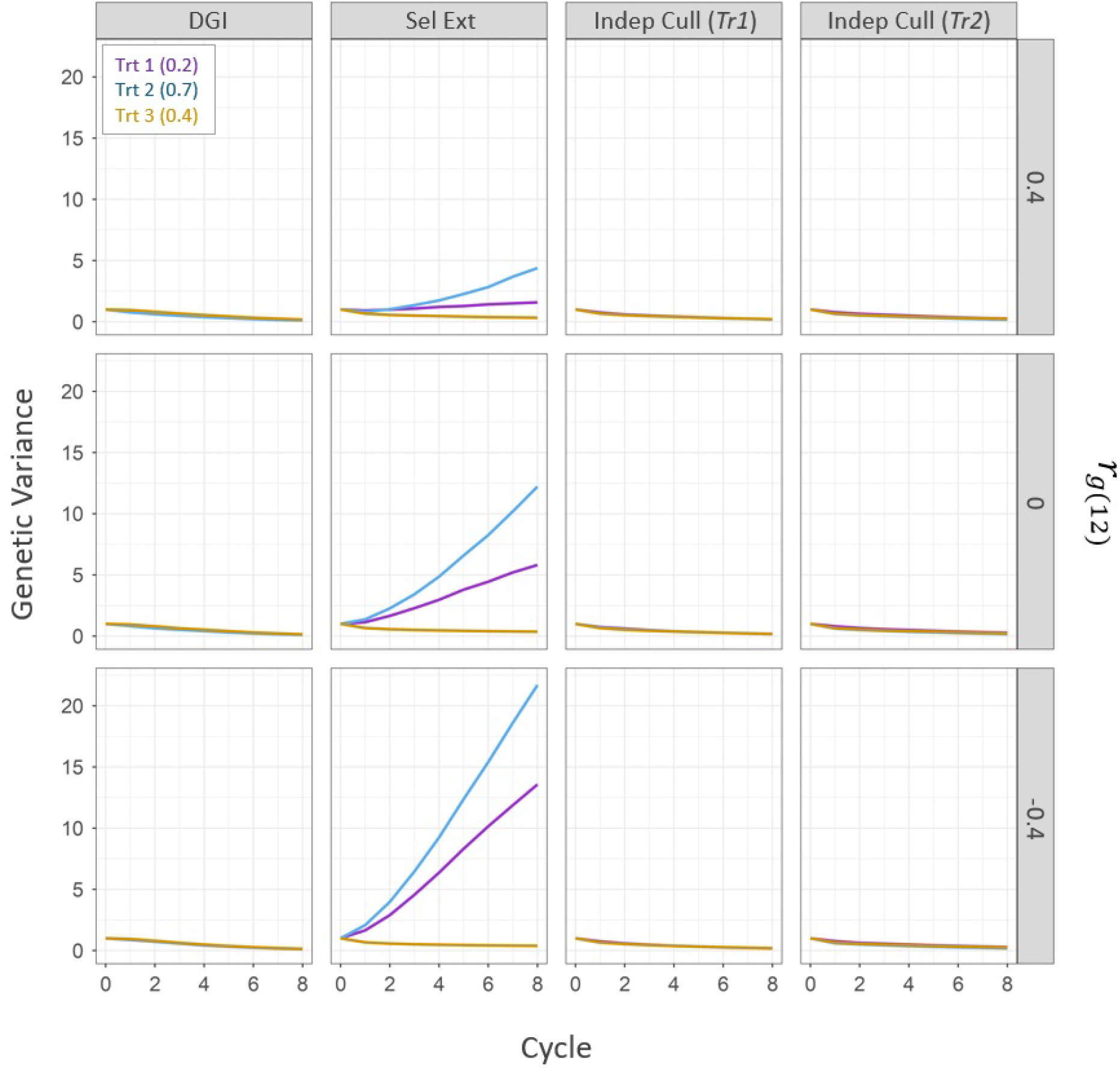
Genetic variance for the three simulated quantitative traits over eight cycles of crossing and parent selection using a Desired Gains Index (DGI), selection of extreme genotypes (Sel Ext), and independent culling (Indep Cull) with genomic estimated breeding values. In each panel, genetic variance is plotted as the change in the mean additive genetic variance over time. Trait 1 is represented by the purple line (Trt 1), trait 2 by the blue line (Trt 2), and trait 3 by the yellow line (Trt 3). Numbers in parentheses indicate the simulated trait heritabilities (*h*^2^) in the founder population. The targeted improvement ratio was 1:1:0 for traits 1, 2, and 3, respectively. For selection of extremes (column 2), initially, trait 3 was stabilised by selecting 30% of the population with values close to zero. Then, the best 15 genotypes were selected for traits 1 and trait 2, respectively (without overlap), and from all possible 225 crosses, 100 were randomly selected to generate the F1. Parent selection using independent culling (columns 3 4) followed a step-wise approach. Initially, trait 3 was stabilised by selecting 30% of the population with values close to zero. Then, the top 10% genotypes for trait 1 were identified from the remaining population, followed by selecting 30 parents based on superior trait 2 values (*Tr1*). Parents were then randomly crossed to generate the F1. To demonstrate the influence of selection order, this process was repeated with trait 2 selection preceding trait 1 (*Tr2*). Each of the selection scenarios (columns) was evaluated under a pairwise genetic correlation (**r**_**g12**_) between trait 1 and trait 2 of 0.4, 0, and -0.4 (rows), respectively. The pairwise genetic correlations between trait 1 and trait 3, and between trait 2 and trait 3, were set to -0.2.

**Fig. S4.**
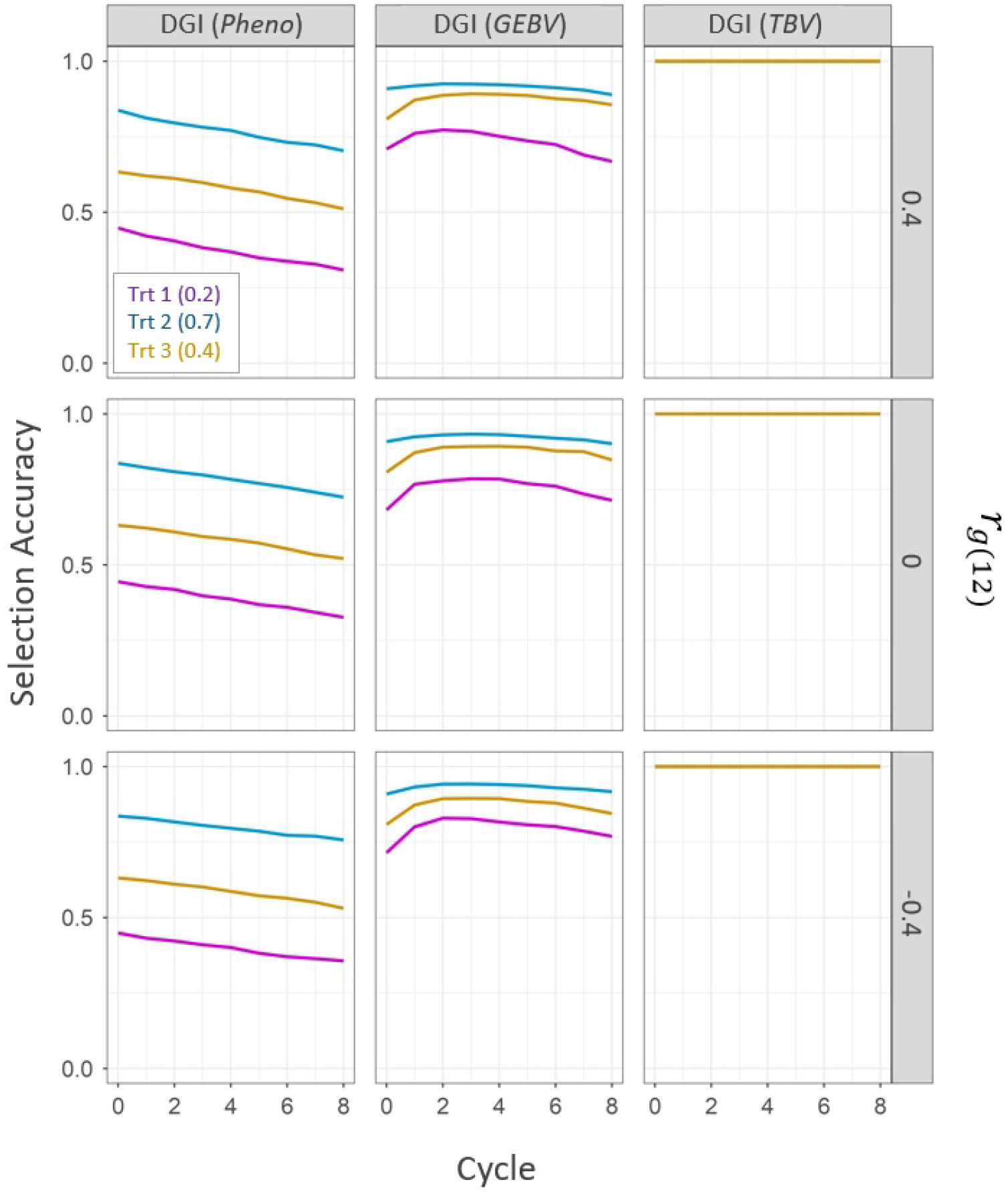
Accuracy of parent selection for the three simulated quantitative traits over eight cycles of crossing and parent selection using a Desired Gains Index (DGI) with: i) phenotypes (pheno), ii) genomic estimated breeding values (GEBV), and iii) simulated true breeding values (TBV). In each panel, selection accuracy is plotted as the change in mean accuracy over time. Trait 1 is represented by the purple line (Trt 1), trait 2 by the blue line (Trt 2), and trait 3 by the yellow line (Trt 3). Numbers in parentheses indicate the simulated trait heritabilities (*h*^2^) in the founder population. Each of the three Desired Gains Index selection scenarios (columns) was evaluated under a pairwise genetic correlation (**r_g12_**) between trait 1 and trait 2 of 0.4, 0, and -0.4 (rows), respectively. The pairwise genetic correlations between trait 1 and trait 3, and between trait 2 and trait 3, were set to -0.2. The breeding objective was to achieve an improvement ratio of 1:1:0 for traits 1, 2, and 3, respectively. Selection accuracy was measured as the Pearson correlation between the true genetic values and the phenotypes or genomic estimated breeding values (GEBVs) of the selection candidates.

**Fig. S5.**
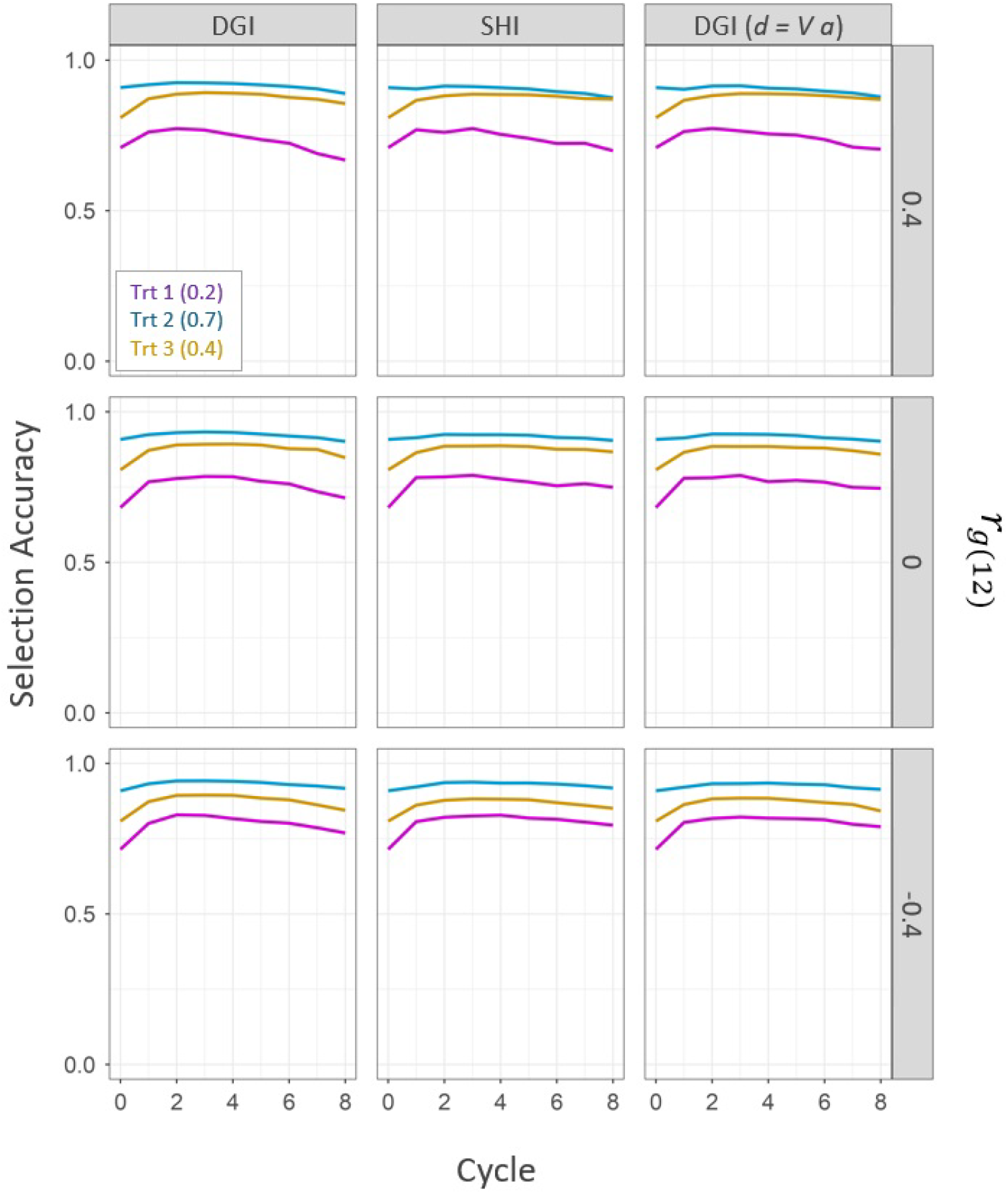
Accuracy of parent selection for the three simulated quantitative traits over eight cycles of crossing and parent selection using a Desired Gains Index (DGI) and a Smith-Hazel Index (SHI) with genomic estimated breeding values. In each panel, selection accuracy is plotted as the change in mean accuracy over time. Trait 1 is represented by the purple line (Trt 1), trait 2 by the blue line (Trt 2), and trait 3 by the yellow line (Trt 3). Numbers in parentheses indicate the simulated trait heritabilities (*h*^2^) in the founder population. For the Desired Gains Index (column 1), the targeted improvement ratio was set to 1:1:0 for traits 1, 2, and 3, respectively. For the Smith-Hazel Index (column 2), the economic weights in vector **a** were set to 1:1:0 for traits 1, 2, and 3, respectively. Column 3 shows a Desired Gains Index with the improvement ratio derived from the vector of economic values **a** (see Eq. 14). Each of the selection scenarios (columns) was evaluated under a pairwise genetic correlation (**r_g12_**) between trait 1 and trait 2 of 0.4, 0, and -0.4 (rows), respectively. The pairwise genetic correlations between trait 1 and trait 3, and between trait 2 and trait 3, were set to -0.2. Selection accuracy was measured as the Pearson correlation between the true genetic values and the phenotypes or genomic estimated breeding values (GEBVs) of the selection candidates.

**Fig. S6.**
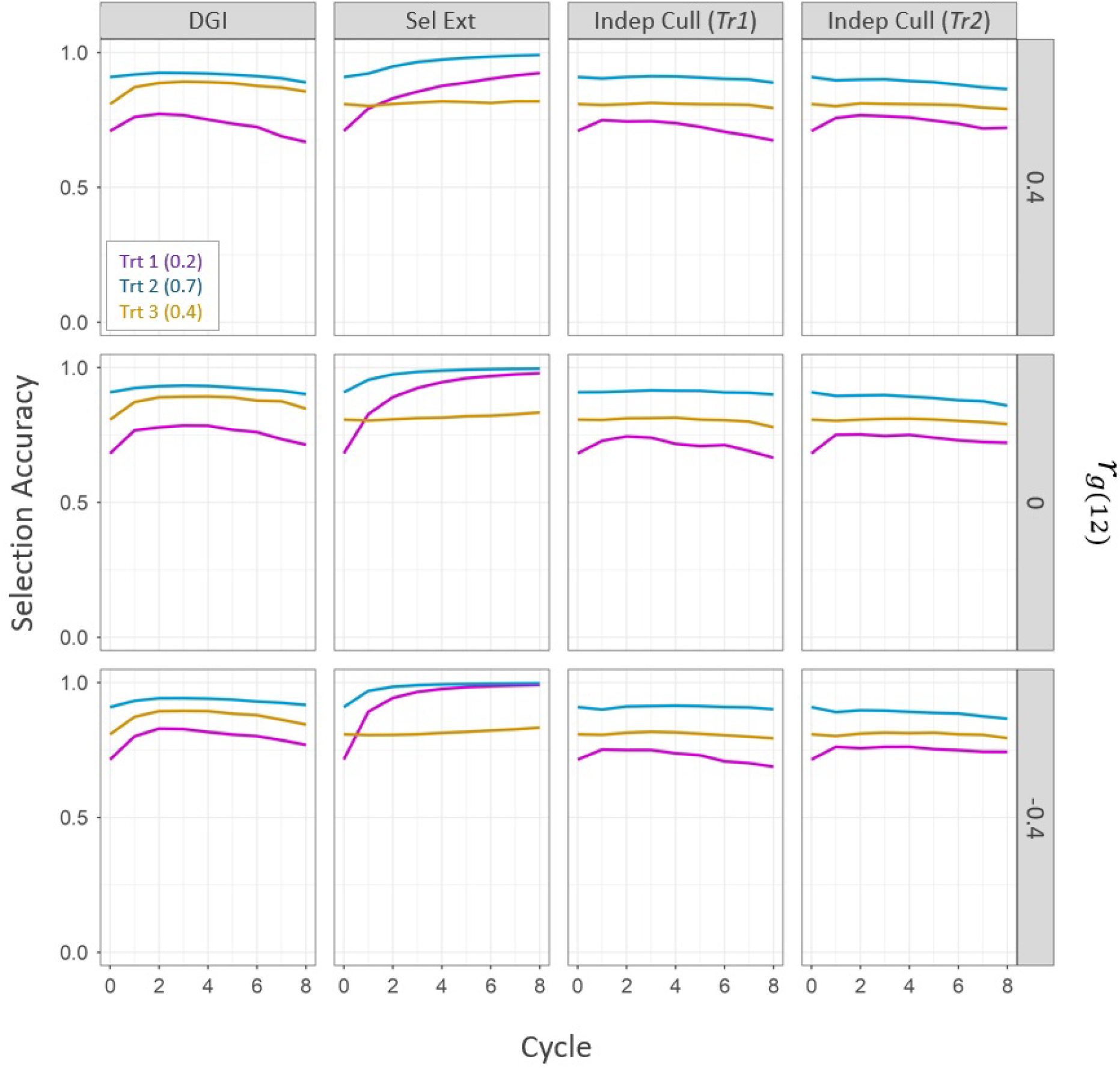
Accuracy of parent selection for the three simulated quantitative traits over eight cycles of crossing and parent selection using a Desired Gains Index (DGI), selection of extreme genotypes (Sel Ext), and independent culling (Indep Cull) with genomic estimated breeding values. In each panel, selection accuracy is plotted as the change in mean accuracy over time. Trait 1 is represented by the purple line (Trt 1), trait 2 by the blue line (Trt 2), and trait 3 by the yellow line (Trt 3). Numbers in parentheses indicate the simulated trait heritabilities (*h*^2^) in the founder population. The targeted improvement ratio was 1:1:0 for traits 1, 2, and 3, respectively. For selection of extremes (column 2), initially, trait 3 was stabilised by selecting 30% of the population with values close to zero. Then, the best 15 genotypes were selected for traits 1 and trait 2, respectively (without overlap), and from all possible 225 crosses, 100 were randomly selected to generate the F1. Parent selection using independent culling (columns 3 4) followed a step-wise approach. Initially, trait 3 was stabilised by selecting 30% of the population with values close to zero. Then, the top 10% genotypes for trait 1 were identified from the remaining population, followed by selecting 30 parents based on superior trait 2 values (*Tr1*). Parents were then randomly crossed to generate the F1. To demonstrate the influence of selection order, this process was repeated with trait 2 selection preceding trait 1 (*Tr2*). Each of the selection scenarios (columns) was evaluated under a pairwise genetic correlation (**r_g12_**) between trait 1 and trait 2 of 0.4, 0, and -0.4 (rows), respectively. The pairwise genetic correlations between trait 1 and trait 3, and between trait 2 and trait 3, were set to -0.2. Selection accuracy was measured as the Pearson correlation between the true genetic values and the phenotypes or genomic estimated breeding values (GEBVs) of the selection candidates.

## References

Baker RJ (1986) Selection Indices in Plant Breeding. CRC Press, DOI 10.1201/9780429280498

Batista LG, Gaynor RC, Margarido GRA, Byrne T, Amer P, Gorjanc G, Hickey JM (2021) Long-term comparison between index selection and optimal independent culling in plant breeding programs with genomic prediction. PLOS ONE 16(5):e0235554, DOI 10.1371/journal.pone.0235554, URL http://dx.doi.org/10.1371/journal.pone.0235554

Bernardo R (2010) Breeding for Quantitative Traits in Plants, 2nd edn. Stemma Press, Woodbury, MN

Brascamp EW (1984) Selection indices with constraints. Animal Breeding Abstracts 52(9):645–65

Bulmer MG (1971) The effect of selection on genetic variability. The American Naturalist 105(943):201–211, DOI 10.1086/282718, URL http://dx.doi.org/10.1086/282718

Chen GK, Marjoram P, Wall JD (2008) Fast and flexible simulation of dna sequence data. Genome Research 19(1):136–142, DOI 10.1101/gr.083634.108, URL http://dx.doi.org/10.1101/gr.083634.108

Covarrubias-Pazaran G, Gebeyehu Z, Gemenet D, Werner C, Labroo M, Sirak S, Coaldrake P, Rabbi I, Kayondo SI, Parkes E, Kanju E, Mbanjo EGN, Agbona A, Kulakow P, Quinn M, Debaene J (2022) Breeding schemes: What are they, how to formalize them, and how to improve them? Frontiers in Plant Science 12, URL https://www.frontiersin.org/journals/plant-science/articles/10.3389/fpls.2021.791859

Erskine W, Williams P, Nakkoul H (1985) Genetic and environmental variation in the seed size, protein, yield, and cooking quality of lentils. Field Crops Research 12:153–161, DOI 10.1016/0378-4290(85)90061-9, URL http://dx.doi.org/10.1016/0378-4290(85)90061-9

Falconer DS, Mackay T (1995) Introduction to Quantitative Genetics, 4th edn. Pearson, Prentice Hall, Harlow, Essex, UK, [16th print]

Finney DJ (1962) Genetic gains under three methods of selection. Genetical Research 3(3):417–423, DOI 10.1017/s0016672300003256, URL http://dx.doi.org/10.1017/S0016672300003256

Gaynor RC, Gorjanc G, Bentley AR, Ober ES, Howell P, Jackson R, Mackay IJ, Hickey JM (2017) A two-part strategy for using genomic selection to develop inbred lines. Crop Science 57(5):2372–2386, DOI 10.2135/cropsci2016.09.0742, URL http://dx.doi.org/10.2135/cropsci2016.09.0742

Gaynor RC, Gorjanc G, Hickey JM (2020) Alphasimr: an r package for breeding program simulations. G3 Genes—Genomes—Genetics 11(2), DOI 10.1093/g3journal/jkaa017, URL http://dx.doi.org/10.1093/g3journal/jkaa017

Gorjanc G, Gaynor RC, Hickey JM (2018) Optimal cross selection for long-term genetic gain in two-part programs with rapid recurrent genomic selection. Theoretical and Applied Genetics 131(9):1953–1966, DOI 10.1007/s00122-018-3125-3, URL http://dx.doi.org/10.1007/s00122-018-3125-3

Hazel L, Dickerson G, Freeman A (1994) The selection index—then, now, and for the future. Journal of Dairy Science 77(10):3236–3251, DOI 10.3168/jds.s0022-0302(94)77265-9, URL http://dx.doi.org/10.3168/jds.S0022-0302(94)77265-9

Hazel LN (1943) The genetic basis for constructing selection indexes. Genetics 28(6):476–490, DOI 10.1093/genetics/28.6.476, URL http://dx.doi.org/10.1093/genetics/28.6.476

Hazel LN, Lush JL (1942) The efficiency of three methods of selection*. Journal of Heredity 33(11):393–399, DOI 10.1093/oxfordjournals.jhered.a105102, URL http://dx.doi.org/10.1093/oxfordjournals. jhered.a105102

Henderson CR (1963) Selection index and expected genetic advance. In: Statistical Genetics in Plant Breeding, vol 982, National Academy of Sciences - National Research Council, Washington D. C., pp 141–163

Henderson CR (1973) Sire evaluation and genetic trends. Journal of Animal Science 1973(Symposium):10–41, DOI 10.1093/ansci/1973.symposium.10, URL http://dx.doi.org/10.1093/ansci/1973.Symposium.10

Hill W (2013) Genetic Correlation, Elsevier, p 237–239. DOI 10.1016/b978-0-12-374984-0.00611-2, URLhttp://dx.doi.org/10.1016/B978-0-12-374984-0.00611-2

Itoh Y, Yamanda Y (1986) Re-examination of selection index for desired gains. Genetics Selection Evolution 18(4):499–504, DOI 10.1186/1297-9686-18-4-499

Kinghorn B (2013) Desire. URL https://bkinghor.une.edu.au/desire.htm, accessed: 2024-07-02

Kwon SH, Torrie JH (1964) Heritability of and interrelationships among traits of two soybean populations1. Crop Science 4(2):196–198, DOI 10.2135/cropsci1964.0011183×000400020023x, URL http://dx.doi.org/10.2135/cropsci1964.0011183×000400020023x

Meredith WR, Bridge RR (1971) Breakup of linkage blocks in cotton, gossypium hirsutum l.1. Crop Science 11(5):695–698, DOI 10.2135/cropsci1971.0011183×001100050027x, URL http://dx.doi.org/10.2135/cropsci1971.0011183×001100050027x

Pešek J, Baker RJ (1969a) Comparison of tandem and index selection in the modified pedigree method of breeding self-pollinated species. Canadian Journal of Plant Science 49(6):773–781, DOI 10.4141/cjps69-132, URL http://dx.doi.org/10.4141/cjps69-13

Pešek J, Baker RJ (1969b) Desired improvement in relation to selection indices. Canadian Journal of Plant Science 49(6):803–804, DOI 10.4141/cjps69-137, URLhttp://dx.doi.org/10.4141/cjps69-137

Powell O, Gaynor RC, Gorjanc G, Werner CR, Hickey JM (2020) A two-part strategy using genomic selection in hybrid crop breeding programs DOI 10.1101/2020.05.24.113258, URL http://dx.doi.org/10.1101/2020.05.24.113258

Smith HF (1936) A discriminant function for plant selection. Annals of Eugenics 7(3):240–250, DOI 10.1111/j.1469-1809.1936.tb02143.x, URL http://dx.doi.org/10.1111/j.1469-1809.1936.tb02143.x

Triboi E, Martre P, Girousse C, Ravel C, Triboi-Blondel AM (2006) Unravelling environmental and genetic relationships between grain yield and nitrogen concentration for wheat. European Journal of Agronomy 25(2):108–118, DOI 10.1016/j.eja.2006.04.004, URL http://dx.doi.org/10.1016/j.eja.2006.04.004

Werner CR, Gaynor RC, Sargent DJ, Lillo A, Gorjanc G, Hickey JM (2023) Genomic selection strategies for clonally propagated crops. Theoretical and Applied Genetics 136(4), DOI 10.1007/s00122-023-04300-6, URL http://dx.doi.org/10.1007/s00122-023-04300-6

Woolliams J, Berg P, Dagnachew B, Meuwissen T (2015) Genetic contributions and their optimization. Journal of Animal Breeding and Genetics 132(2):89–99, DOI 10.1111/jbg.12148, URL http://dx.doi.org/10.1111/jbg.12148

Yamada Y, Yokouchi K, Nishida A (1975) Selection index when genetic gains of individual traits are of primary concern. The Japanese journal of genetics 50:33– 41, URL http://dx.doi.org/10.1266/jjg.50.33

Young SSY (1961) A further examination of the relative efficiency of three methods of selection for genetic gains under less-restricted conditions. Genetical Research 2(1):106–121, DOI 10.1017/s0016672300000598, URL http://dx.doi.org/10.1017/S0016672300000598

